# Assessing extracellular vesicles in human biofluids using flow-based analyzers

**DOI:** 10.1101/2022.07.20.500853

**Authors:** Olga Krzyzaniak, Kevin Ho Wai Yim, Ala’a Al Hrout, Ben Peacock, Richard Chahwan

**Affiliations:** Institute of Experimental Immunology, University of Zurich, 8057 Zurich, CH; NanoFCM, ltd. Nottingham, UK

**Keywords:** extracellular vesicles, nano-analyzer, nanoFCM, liquid biopsy, biofluids

## Abstract

Extracellular vesicles (EVs) are increasingly being analyzed by flow cytometry. Yet, their miniscule size and low refractive index, causes the scatter intensity of most EVs to fall below the detection limit of most flow cytometers. A new class of devices, known as spectral flow analyzers, are becoming standards in cell phenotyping studies. Largely, due to their unique capacity of detecting a vast panel of markers with higher sensitivity for light scatter detection. Another class of devices, known as nano-analyzers, provides high resolution detection of sub-micron sized particles. Here, we aim to compare the EVs phenotyping performance between the Aurora (Cytek) spectral cell analyzer and the NanoFCM (nFCM) nanoflow analyzer. These two devices were specifically chosen given their lead in becoming gold standards in their respective fields. Immune cell-derived EVs remain poorly characterized despite their clinical potentials. We therefore, used B- and T- cell line-derived EVs and donor-matched human biofluid-derived EVs from serum, urine, and saliva in combination with a panel of established immune markers for this comparative study. A comparative evaluation of both cytometry platforms was performed, discussing their potential and suitability for different applications. We found that nFCM can accurately i) analyze small EVs (40 to 200 nm) matching the size accuracy of electron microscopy; ii) measure concentration of single EV particle per volume; iii) identify underrepresented EV marker subsets; and iv) provide co-localization of EV surface markers. We could also show that human sample biofluids have unique EV marker signatures that could have future clinical relevance.

## INTRODUCTION

Extracellular vesicles (EVs) are nano to micro-sized (40-1000nm) lipid bilayer vesicles released by all types of cells for intercellular communication through a plethora of signaling modalities. EV cargo molecules include nucleic acids, proteins, lipids, and other bio-molecules(1, 2). EVs are found in all biological fluids and recent studies have shown their promising potential in disease diagnosis and therapeutics as well as vaccine development (1, 3–6). However, due to their small size and low refractive index, the scatter intensity of most EVs is below the detection limit of common flow cytometers (7, 8). These hurdles have significantly slowed down the research progress of understanding the role of EVs in different life science research fields.

The active involvement of EVs in immune signaling and regulatory processes(1), ushers an urgent need for reliable and efficient turn-over analytics for immune EVs as a quality assessment, as well as a novel immune phenotyping, tools. Flow cytometry-based EV analyses mostly rely on antibody-bound bead based capturing methods (9, 10), which help to overcome the size detection limit of conventional cytometers. Unfortunately, as no universally agreed upon EV marker is currently known (11–13), this approach has the crucial drawback of overlooking all EV populations which do not express the specific capture marker of interest (5). In addition, this type of analysis cannot provide single particle resolution since one bead can bind several EVs simultaneously and vise-versa.

Label-free single particle nanoflow analyzers, first brought by NanoFCM (nFCM) nanoflow analyzer, provide the rare possibility of phenotyping EVs with the least pre-analytical filtering steps and the results were proven by several studies in the past years(3, 14). Due to its physical design however, the number of fluorophore choices and their combinations are limited in the nFCM by a two-laser conformation. Hence, we attempted to compare this system to a new class of flow cytometer, namely Aurora (Cytek) spectral flow analyzer which utilizes photomultiplier tubes and a high number of detectors, allowing it to increase the number of resolvable fluorophores as well as its overall sensitivity in particles detection. Since immunological studies mostly focus on cell-to-cell interactions and signaling in the circulation of biofluids (15–17), especially peripheral blood, we first benchmarked the two platforms using B- and T-cell line derived EVs to verify the proposed antibodies panel and flow cytometers configuration(18). The verified panel was further applied to analyze biofluid-derived EVs to reproduce the conditions of immunological and clinical studies through easily accessible liquid biopsies. We compared the size, concentration, surface marker quantifications of sample EVs between the two platforms and ultimately describe the beneficial properties of each platform for varying application goals. We also attempt to identify unique EV signatures in healthy biofluid donors to extrapolate clinical potentials.

## MATERIALS AND METHODS

### Cell cultures

Jurkat and Ramos cells were maintained in RPMI-1640 supplemented with 10% FCS, 1% Pen-Strep, 1% Pyruvate, 5% NCTC-109, and 0.1% B-Mercaptoethanol and split once 90% confluency was reached. Cells were cultured in EV depleted FCS media (achieved by 100 kDA ultrafiltration column, 3,000 g for 55 mins) at 10^7^ cells/mL overnight before harvesting EVs. Conditioned media were centrifuged at 300 g to remove cells, 2,000 g to remove debris, 10,000 g to remove large particles, and the remaining conditioned media were subjected to 100,000 g for final purification.

### Biofluids processing

Peripheral blood, urine, and saliva were drawn from healthy volunteers. For blood EVs, 3 – 5 mL of peripheral venous blood was collected from into BD Vacutainer serum clot activator 10 ml (367896). After collection, tubes were left vertically undisturbed on the bench for 15 minutes to allow blood clot and followed by centrifugation at 2,500 g for 10 mins at 4°C for separation of sera. Sera were centrifuged at 10,000 g for 30 mins at 4°C to remove large particles. Clarified serum was split into 0.5mL aliquots and stored in -80°C until further use. 1 mL of saliva was diluted in 4 mL of 0.22 µM filtered PBS. 3-5 mL of urine and diluted saliva were centrifuged at 2,500 g for 10 mins at 4°C and the remaining supernatants were centrifuged at 10,000 g for g for 30 mins at 4°C to remove large particles. Clarified urine and saliva supernatant was stored in 0.5 mL and 0.15 mL aliquots respectively and stored in -80°C until further use.

### Isolation and immunostaining and nanoflow analysis of purified EVs

Frozen biofluid EV aliquots were thawed quickly at 37°C and diluted with 5x volume of 0.22 µM filtered PBS. 1 mL of 10,000 g spun conditioned medium from Ramos and Jurkat cells, and 1 mL of diluted biofluids samples were used for each staining panel. Samples were centrifuged at 100,000 g and 4°C for 1 hour. Purified EVs were stained in 100 µL staining buffer with antibodies listed in table S1 for 30 mins in room temperature covered in dark. Following the incubation, select samples were stained with 1uL of Celltrace in DMSO (1:10 working solution) and incubated in dark for a further 30min. Stained EVs were washed with 1 mL 0.22 µM filtered PBS at 100,000 g for 45 mins. Washed EVs were resuspended in 50 µl of 0.22 µm filtered PBS and subjected to flow analysis by NanoFCM Flow Nanoanalyzer. Following this analysis, the samples were further diluted, to minimize abort rate due to swarming, and analyzed with the Aurora. For purity assessment, triton-X was added to purified EVs at final concentration of 1% followed by vigorous vortex for approximately 30 seconds before acquisition.

### Purified EVs analysis with nFCM

Monodisperse silica nanoparticles of four different sizes, with modal peak sizes of 68 nm, 91 nm, 113 nm and 155 nm were used as the size reference standard to calibrate the size distribution of EVs. Samples were recorded for 1 minute with the range of 2,000 to 12,000 events per minute to minimize the swarm detection effect since there is enough time/space between each particle measurement as evinced by the event burst trace which shows separate measurements of particles. The steady flow of the system allows for comparison of particle detection rate to a concentration standard (a stable 250nm silica bead of 1.87e10/ml (batch variable)). The particle concentration is then calculated, including the dilution factor, in particles/ml.

### Purified EVs analysis with Aurora

Threshold for side scatter was set to 800 and gain of B2 (FITC) and R1 (APC) detectors were set to 2500. Apogee sizing beads (#1527) with mixture of beads ranging from 80 to 1300 nm were used as the size reference to confirm the particle resolution of the analyzer. PBS was used as blank and sample acquisition was performed until event rate of PBS reduced to 1000 events per second. Unless stated otherwise, for all samples 80,000 events were recorded with the slowest flow rate (10 to 15 µl per second) to reduce swarm effect.

### Data and statistical analysis

EV flow cytometry data was exported as FCS files and analyzed using Flowjo software (Treestar). Statistical analysis of flow cytometry values was performed using Graphpad (version 9.1.1, GraphPad Software, La Jolla California USA). Ordinary One-way ANOVA, P-value < 0.05 was considered as statistical significance. PCA and biplot were generated with R software (version 4.0.1) using the package ‘factoextra’ and ‘ggplot2’. Spearman’s correlation analyses were produced using the corrplot package and implemented hclust clustering method. Hierarchical clustering in the heatmap was performed using the Euclidean distance, through the Morpheus online software (Morpheus Broad Institute, https://software.broadinstitute.org/morpheus). The correlations between different EVs subsets were analyzed using non-parametric Spearman correlations. The significance threshold was set at alpha < 0.05.

### Transmission electron microscopy of purified EVs

To visualize EVs samples under the transmission electron microscope samples were transferred onto pioloform-coated EM copper grids by floating the grids on a droplet containing freshly prepared exosome placed on parafilm. After 5 min of incubation the grids were washed 3 × 5 min on droplets of deionized water before contrasting of bound exosomes in a mixture of 2% w/v methyl cellulose and 2% w/v uranyl acetate (mix 9:1) on ice for 10 min. Contrasted grids were then air dried in a wire loop before analysis using a FEI Talos 120v. Images were taken with digital CMOS camera BM-Ceta (Thermo Fisher). High-resolution images and particle size quantification analysis were acquired and performed by MAPS software (Thermo Fisher).

## RESULTS AND DISCUSSION

### Comparison of properties and sizing capacities in cell-line derived EVs

To compare the EV phenotyping capacity (size, concentration, and surface marker profile) between Aurora and nFCM, we started by benchmarking cell-line derived EVs, because of their i) size homogeneity, ii) controlled and reproducible yield from *in vitro* culture production, and iii) conserved parental cell marker expression for antibody panel benchmarking. Cells from Jurkat, a human T cell leukemia cell-line, and Ramos, a Burkitt’s lymphoma B-cell line, were seeded in EV depleted FCS media overnight at the cell density of 10^7^/mL, after which EVs were harvested by serial centrifugation before flow analysis. The same sample sets were then sequentially measured on both devices to compare their phenotyping capabilities as accurately as possible. Size distribution of both Jurkat and Ramos EVs was visualized using side scatter histograms (Fig. 1A). Vesicle size distribution was determined using reference beads from Apogee (default beads for Aurora analysis) and nFCM’s four-peak beads (default beads for nFCM) with separate gates spanning the scatter signals area underneath each sizing bead populations to enhance quantification power of sample vesicle size distribution (Fig. S1). The vast majority of purified EVs from both cell lines were below 180nm and even below 68 nm – the smallest reference-bead population sizes of the Aurora and nFCM respectively (Fig. 1A). This suggests that the EV purification protocol is efficient for isolation of small EVs (sEVs; 40-150nm) in particular. Furthermore, since the Aurora has a resolution threshold of approximately 100nm, this indicates that the instrument was unable to further resolve the particle size distribution by binning according to scatter peak signals of size reference bead scatter. Purity of isolated EVs were assessed by 0.1% Triton-X treatment, which is known to disrupt EV integrity, with around 80% reduction in particle counts and Celltrace Far Red signals, respectively (Fig. S2). In the case of nFCM, Jurkat EVs were slightly, but not significantly, more enriched in the 68nm peak gate compared to Ramos EVs, while the latter was marginally more enriched in 91 nm, 113 nm, and 155 nm peak gates compared Jurkat EVs (Fig. 1B). The same size binning could not be achieved in Aurora due to low size resolution for sEVs and the background noise we observed in PBS controls at that size interval (Fig. 1A). However, we could identify larger EV size bins and for which both cell lines seemed indistinguishable from each other (Fig. 1B). Furthermore, using the nFCM size conversion algorithm based on the standard curve generated by sizing beads data, the overall mean size of sample EVs were determined (Fig. 1C).

**Figure 1.**
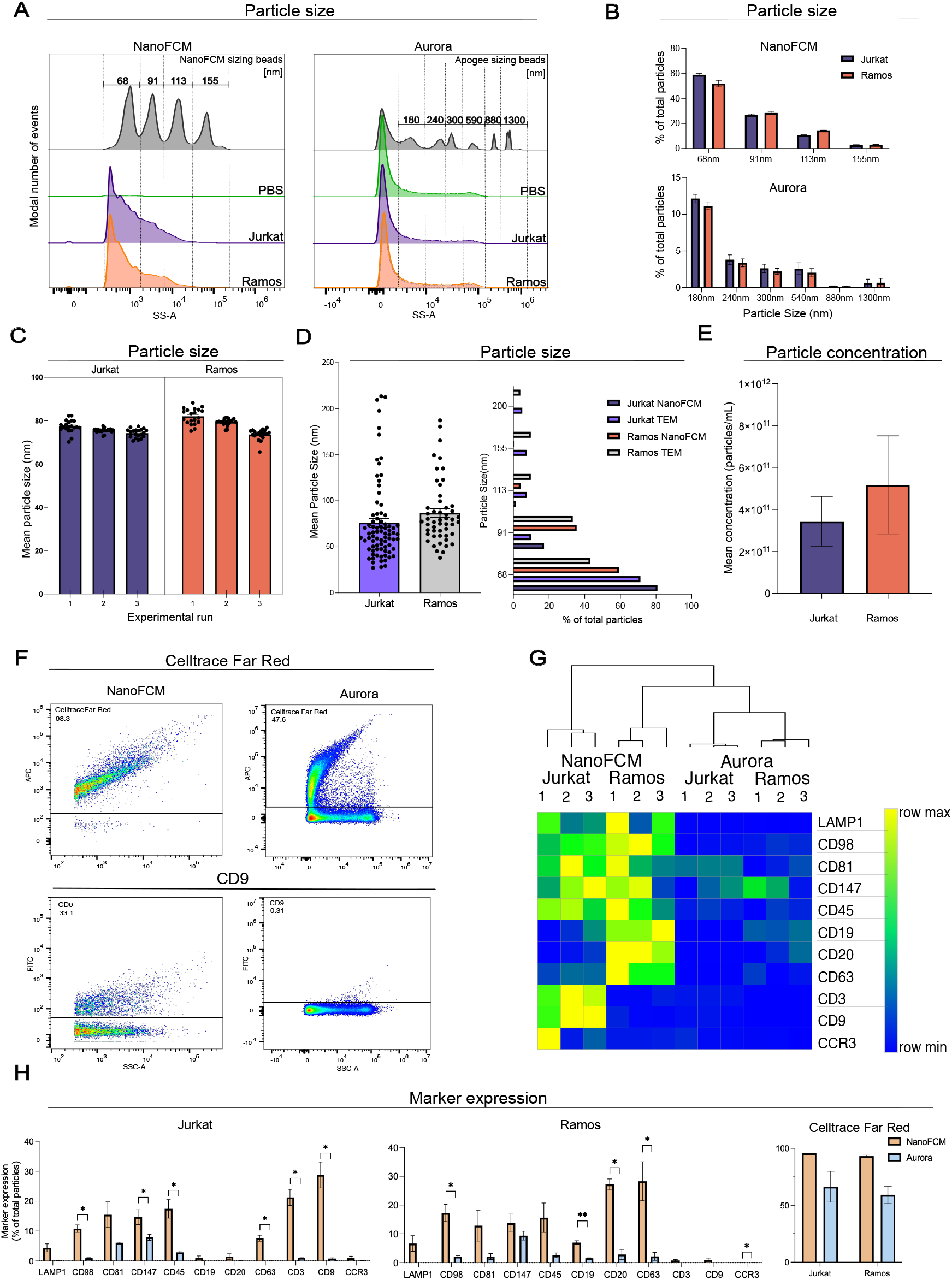
Jurkat and Ramos derived sEVs phenotyping using Cytek^®^ Aurora and NanoFCM^®^ nanoflow analyzer. (A) Representative scatter histograms of comparison between the particle size distribution of Jurkat and Ramos derived sEVs with nFCM sizing beads (68, 91, 113, 155 nm) (left, nFCM) and Apogee sizing beads (right, Aurora), scatter area under modal peaks of sizing beads were used to gate for different particle size intervals. (B) Approximated size distribution quantification of Jurkat and Ramos derived sEVs measured in nFCM and Aurora with particle size binning strategy in (A). (C) Mean approximated particle size of sEVs in each nFCM experimental run as calculated by the nFCM analysis software. (D) Approximated size distribution and mean particle size quantification of Jurkat and Ramos derived sEVs TEM compared to nFCM analysis using particle size binning strategy in (A). (E) Particle concentration quantification of Jurkat and Ramos derived sEVs measured in nFCM using standard QC beads with known concentration. (F) Representative dot plots of CellTrace Far Red (R1 in Aurora; PC5 in NanoFCM) and CD81 (B2 in Aurora; B2 in NanoFCM) to demonstrate gating strategy. (G) Clustered heatmap of denoted sEVs markers expression of Jurkat and Ramos sEVs measured by nFCM and Aurora using Euclidean distance calculation. (H) Quantification of sEVs markers of Jurkat and Ramos derived sEVs measured by nFCM and Aurora, values are representing percentage of total measured particles. Multiple T-test, *p* < 0.05 *, *p* < 0.01 **, *p* < 0.005 ***, n=3.

### EV sizing by nFCM highly reproduces electron microscopy standards

Mean EV sizes generated in nFCM were then compared to the transmission electron microscopy (TEM) which remains the *de facto* gold standard for accurate EV sizing (Fig. 1D). With the aid of MAPS high resolution image analysis software, mean diameter of Jurkat and Ramos cell EVs were determined from over 80 and 50 high resolution TEM imaged EVs, respectively. Compared to size distribution intervals data from nFCM, both Jurkat and Ramos cell EVs displayed very similar distribution profile of EV sizes, suggesting the sizing performance of nFCM is very comparable to the TEM; hence accurate, easier to measure, and amenable to high throughput analyses. Additionally, given the capacity of quantifying burst traces of sample particles in nFCM, thereby providing higher control in avoiding swarm effects, the software can produce direct particle concentration conversion using QC beads of known concentration. Particle concentration yielded from same number of EVs producing cells could therefore be quantified for the nFCM collected data (Fig. 1E).

### Comparison of phenotyping capacities in cell-line derived EVs

Next, we compared the fluorescence detection performance of selected enriched EVs markers, immune markers, and membrane dye of EVs from both cell-lines using the two different platforms (Fig. 1F). Our antibody panel consisted of recently identified exosome-specific LAMP1, ectosome-specific CD147 (BSG) and CD98 (SLC3A2), cell membrane dye (CellTrace Far Red), tetraspanins (CD9, CD63, and CD81), pan-leukocyte CD45, pan-B cell CD19 and CD20, pan-T cell CD3, natural killer specific CD56, classic monocytic CD14, granulocytes specific CD66b and CCR3. Isotype control antibodies IgG1 as well as IgG2a (both rat and mouse) were added as negative controls, in addition to unstained, antibody in PBS only, PBS only and TritonX detergent treated samples (Fig S3). As expected, pan-B cell markers CD19 and CD20 expression was below 2% in Jurkat EVs, while the pan-T cell marker CD3 expression was below 2% in Ramos EVs measured in both platforms confirming the homogeneity of purified EVs and the specificities of antibodies (Fig. 1G). CellTrace signals were predominantly over 90% in all samples measured with the nFCM platform, confirming the purity of EVs analyzed. CellTrace signal was strongly reduced when the same samples were measured in parallel on the Aurora (Fig. 1H). Interestingly, neutrophil specific CD66b was observed in rather high percentage of Jurkat and Ramos EVs in nFCM at 12.11% and 4.46% respectively. In a recent RNA-seq study investigating immune cells RNA signature, transcript expression of CD66b was higher in several B cell subsets than neutrophils(19). Previous cell studies have shown CD66a, in the same family as CD66b, is expressed on B and T cells suggesting lymphocytes may also encode CD66b, which is then being shuttled by secreted EVs instead of expressing on their cell surface (20, 21). In Jurkat EVs measured by nFCM, the top 3 expressed markers were CD81, CD3, and CD147 at 25.60%, 23.80%, and 17% of total particles, respectively. This is relatively similar though not identical to results from Aurora, where the top 3 markers were CD81, CD147, and CD45 at 19.65%, 7.53%, and 3.20% of total particles, respectively. In Ramos EVs measured by nFCM, the top 3 expressed markers were CD20, CD63, and CD81 at 26.20%, 19.95%, and 18.90% of total particles respectively. Meanwhile in Aurora, the top 3 markers were CD81, CD63, and CD147 at 15.45%, 13.15% and 8.27% respectively. Although the relative EV marker expression within the given panel was similar between Aurora and NanoFCM in a given source of EVs, the absolute percentages of such EV subsets were very different between the two platforms, e.g. 12% vs 1.15% CD3 in Jurkat EVs and 47.8% vs 16.7% CD20 in Ramos EVs. This can possibly be due to differences in background noise level and particle size resolving window between the platforms.

### Comparison of phenotyping capacities in three human biofluids derived EVs

After establishing the fundamental parameters using cell-line EVs, we proceeded with analysis of more complex biological biofluids. Serum, urine and saliva derived sEVs were used for phenotyping comparison due to their relevance in clinical studies and diagnostics. Size distributions of different biofluid derived EVs were visualized using side scatter histograms (Fig. 2A). Using size reference beads from Apogee (for Aurora analysis) and nFCM (four peaks beads), we determined the majority of purified EVs in our samples to be below 180 nm, as measured in the Aurora, and more specifically below 155 nm as measured in nFCM. Scatter histograms from both Aurora and nFCM showed larger vesicles were presented in serum EVs compared to urine and saliva EVs (Fig. 2A). nFCM analysis was able to further resolve the differences in approximate size distribution between different sources of EVs, with saliva and urine EVs being more enriched than serum in the 68nm bin, while the latter was more enriched in bins of 91nm and above. Mean particle size data generated by the nFCM software reports also supported this result, showing serum vesicles to be on average larger, with a mean particle size of 73.6nm, then urine and saliva vesicles, which had mean particle sizes of 67.4nm and 65.3nm respectively (Fig. 2B). Interestingly, mean sEV concentration was found to be lowest in serum with 1.5×10^10^ particles/mL, followed by urine with 1.1×10^11^ particles/mL and finally saliva with the highest concentration of EVs at 4.84×10^12^ particles/mL (Fig. 2C). Purity of isolated EVs were also assessed by Celltrace Far Red staining (Fig. 2D). Treatment with 1% Triton-X detergent, indicated high sample purity in all biofluids where mean concentration reduction in serum, urine and saliva was determined to be 84%, 85% and 84% respectively (Fig. S4).

**Figure 2.**
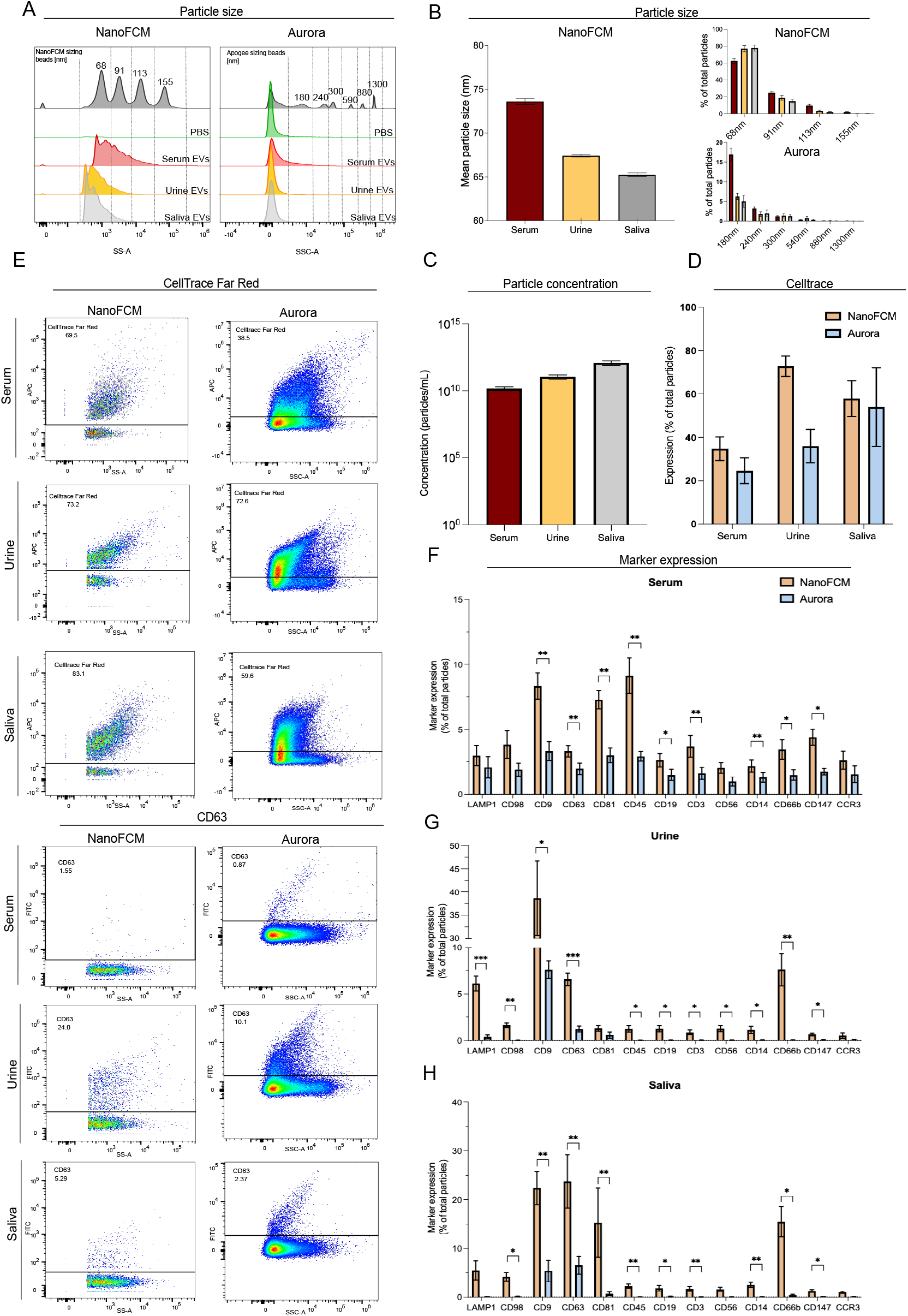
Healthy human donor biofluids derived sEVs phenotyping using Cytek® Aurora and NanoFCM® nanoflow analyzer. (A) Representative scatter histograms of comparison between the particle size distribution of serum, urine, and saliva derived sEVs with nFCM sizing beads (68, 91, 113, 155 nm) (left, nFCM) and Apogee sizing beads (right, Aurora), scatter area under modal peaks of sizing beads were used to gate for different particle size intervals. (B) Mean approximated particle size of denoted sEVs calculated by the nFCM analysis software (left). Approximated size distribution quantification of denoted sEVs measured in nFCM and Aurora with particle size binning strategy in (1A) (right). (C) Particle concentration quantification of denoted biofluids derived sEVs measured in nFCM using standard QC beads with known concentration. (D) Quantification of CellTrace Far Red intensity in denoted biofluids derived sEVs measured by nFCM and Aurora for EVs purity assessment. (E) Representative dot plots of CellTrace Far Red (R1 in Aurora; PC5 in nFCM) and CD63 (B2 in Aurora; B2 in nFCM) in denoted sEVs to demonstrate gating strategy. (F-H) Quantification of sEVs markers of serum (F), saliva (G), and urine (H) derived sEVs measured by nFCM and Aurora, values are representing percentage of total measured particles. Multiple T-test, *p* < 0.05 *, *p* < 0.01 **, *p* < 0.005 ***, n=4.

Fluorescence detection performance of selected enriched immune EVs markers and membrane dye of biofluid derived EVs was also compared between the two different platforms (Fig. 2E-H). Isotype controls were performed to confirm specificities of antibodies used (Fig. S5). In serum EVs, the top four markers detected in nFCM were CD45, CD9, CD81 and CD147 and similarly CD45, CD9, CD81 and CD63 in Aurora, which fits well with expectations for this type of biofluid. In urine EVs, CD9 was the most expressed marker measured in both platforms, however CD66b, CD63 and LAMP1, which were detected in 7.6%, 6.6% and 6.1% of particles by nFCM, could not be detected by Aurora. Similarly, in saliva sEVs, CD63 and CD9 were readily detected to different extents by both platforms, with CD63 expression being detected in 23.7% of particles in nFCM and 6.5% in Aurora and CD9 expression in 22.4% in nFCM and 5.4% in Aurora. The less expressed CD81 and CD66b could only be detected by nFCM however, where they were detected in 15.5% and 15.3% of vesicles respectively, and almost not at all in Aurora. In general, specialized immune cell markers, CD19, CD3, CD56, CD14, CD66b, and CCR3 were more enriched in serum EVs compared to urine and saliva, most likely due to the predominant localization and intercellular signaling between blood cells. Comparatively speaking, the two platforms showed similar relative EVs markers expressions within the given panel, however, the absolute percentages of markers between the two platforms were very different, 50% vs 10% CD9 in urine EVs, most likely due to background noise level differences and particle size resolving window between the platforms. Moreover, lowly expressed markers were often below 1% positive in Aurora, making *de novo* EVs markers identification more challenging compared to nFCM.

### EVs markers can define the origin of biofluids irrespective of the donor

Next, we attempted to define the origin of sEVs from different biofluids according to the distribution of our panel of selected markers. We performed principal component analysis (PCA) and dimension reduction correlation analysis of EVs markers derived from different biofluids using the values from nFCM measurement due to its superiority in sEVs markers detection seen in above sections. From the unsupervised PCA Bi-plot exploratory analysis of four independent donors derived serum, urine, and saliva sEVs markers, clear stratification of EVs origin were observed regardless of donor’s variabilities (Fig. 3A). Interestingly, CD3 (T-cell specific), CD45 (pan-leukocytes), and CD147 (plasma membrane protein) are the strongest drivers for the stratification between serum and other EVs. Urine and saliva EVs signatures were not as obvious as serum EVs, with CD9 and LAMP1 (exosomes enriched) being the strongest drivers for urine derived EVs. CD66b (granulocytes specific) and CD63 were more represented in saliva EVs, while specialized immune EVs expressing CD14 (classical monocytes), CD19 (B-cell specific), CD56 (Natural Killer-cell specific), and CD98 (ectosomes enriched) markers were similarly represented in serum and saliva EVs, that drove stratification from urine EVs. Spearman’s correlation analysis of EVs markers from different biofluids origins also revealed strong positive correlations between specialized immune EVs derived from serum; between multiple markers in saliva EVs and urine EVs. Strong negative correlations were also observed between markers from serum EVs and urine EVs (Fig. 3B). Bi-plot and Spearman’s correlation analysis in each type of biofluids measured in both Aurora and nFCM also revealed very different PC loadings and correlation patterns between our selected EVs markers, suggesting the EVs landscape varied significantly in different biofluids (Fig. S6). These results further highlight the differential power of flow-based EVs markers analysis in identification of the origin of certain EVs subsets, which is highly relevant in the diagnostic and prognostic application in clinical settings.

**Figure 3.**
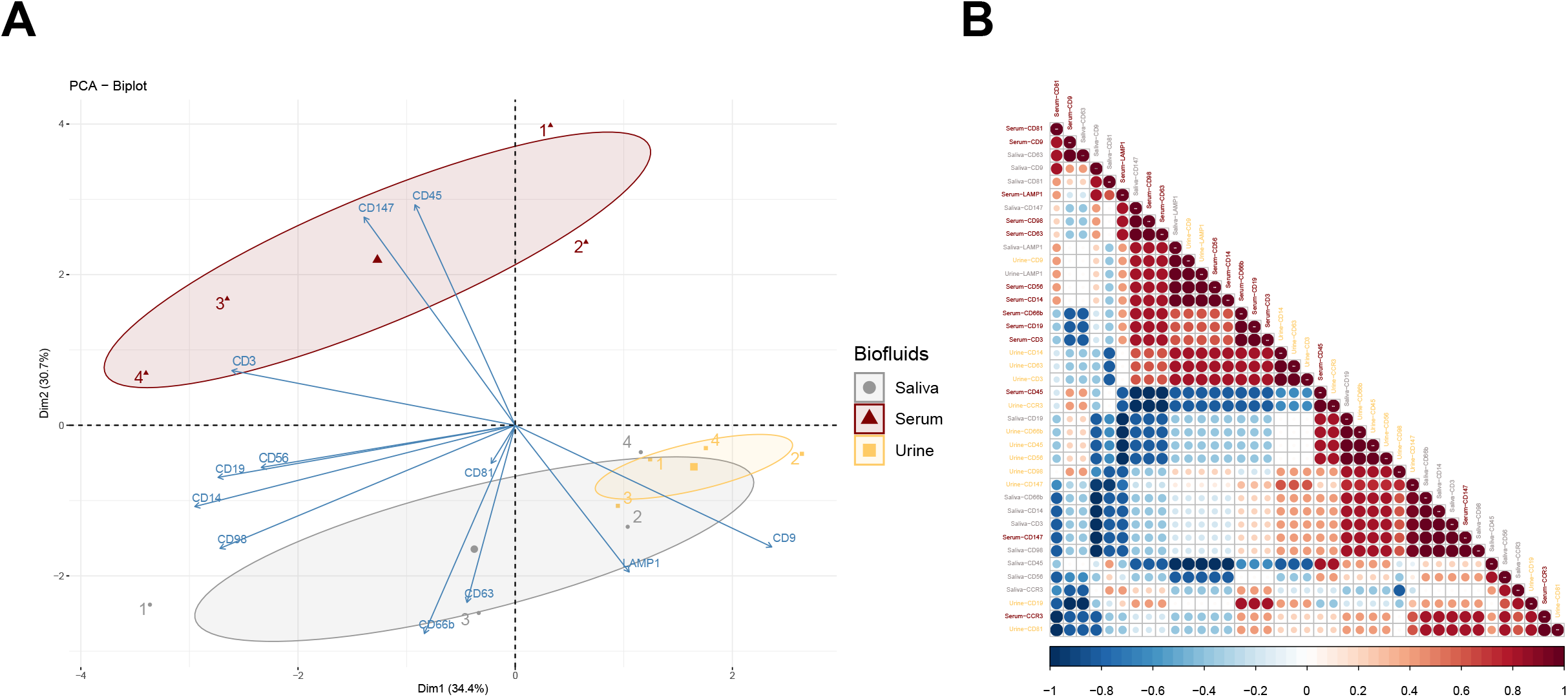
PCA and correlation of biofluids derived sEVs markers measured in NanoFCM® nanoflow analyzer. (A) PCA bi-plot analysis of EVs markers measured by NanoFCM stratifying different sources of biofluids EVs. Four individual human donors indicated as 1, 2, 3, and 4. (B) Spearman’s rank correlation matrix of serum, urine and saliva sEVs markers from data acquired by NanoFCM. N=4. One-way ANOVA, p < 0.05 *, p < 0.01 **, p < 0.005 ***.

### Comparative evaluation of Aurora versus nFCM flow cytometers for EV analyses

The swarming effect of the EVs is a further factor which should be taken into consideration when deciding between the two systems. While the Aurora does offer the option of lowering flow rate, allowing us to reduce the swarming, this is not enough to alleviate the issue entirely. The abort rate throughout the samples was consistently high and diluting them only helped to resolve the issue marginally. Furthermore, we noticed some of the Auroras we tested were able to avoid swarming more efficiently than others, indicating that instrument set-up can have a significant effect on the final experiment outcomes. NFCM on the other hand was able to reduce the swarming effectively, allowing us not only to retain a larger portion of the signal but also to effectively quantify the vesicles based on side scatter.

Depending on research aims and priorities, there are benefits and considerations between the two platforms for EVs studies. Our concluding results can be aptly summarized as shown in Table 1. Since Aurora has been designed as a flow cytometer for cell phenotyping, it has a wider particle size window of sample analysis compared to nFCM which is specifically designed for sub-micron particles. The former platform would be useful if the samples are more heterogenous and have a bigger size range. nFCM would be more beneficial if the samples are known to be sized under 200 nm. In terms of concentration quantification, the ability of quantifying burst trace signals from scattering of each particle in nFCM provides a surplus advantage in phenotyping analysis as well as accurate EVs loading control for functional assays. In the context of surface marker profiling, nFCM is less affected by background noise possibly coming from photodiode detectors and sheath flow in conventional cytometers. Hence, EVs fluorescence signals are less masked by the high background noise which is usually mixed within the negative EVs population and ultimately leads to more accurate readout and increased sensitivity especially in the case of detecting lowly expressed markers compared to the Aurora. In terms of fluorophore choices, the Aurora is equipped with five lasers and able to detect up to 40 different types of fluorophores, while nFCM has only two lasers and a much tighter emission detection bandwidth. Provided the marker of interest is highly expressed and the samples are in the > 100 nm size range, one could streamline their staining panel based on the flexible fluorophore options in Aurora, to simultaneously co-stain at least two or three markers of interest with the furthest emission wavelength from each to avoid spillover. Lastly, in our hands, the Aurora requires less time and expertise for calibration, operation, and data analysis compared to nFCM, which are elements to be considered in mass scale studies and high-throughput settings.

**Table 1.** Comparative table of benefits and considerations in Cytek® Aurora and NanoFCM® nanoflow analyzer for EVs analysis.

| Parameters | Platforms | Cytek Aurora | NanoFCM |
| --- | --- | --- | --- |
| Resolvable particle size |  | > 100 nm | 40 - 200 nm |
| Particle concentration estimation |  | ↓ (higher swarm effect) | ↑ (Less swarm effect/Particle visualized as direct burst signals) |
| Side scatter detection of sEVs (< 100 nm) |  | ↓ (higher overlap with noise) | ↑ (less overlap with noise) |
| Fluorescence detection of labeled sEVs (< 100 nm) |  | ↓ (Signals weaken by higher noise) | ↑ (Less noise) |
| Fluorophore choices |  | ↑ (5 lasers & up to 40 colors) | ↓ (2 lasers & tighter emission bandwidth detection) |
| Ease of Operation & Throughput |  | ↑ (Low expertise required set-up & < 1 min/sample) | ↓ (Expertise required set-up & ~ 2 mins/sample) |

## ACKNOWLEDGEMENTS AND FUNDING

RC is supported by BBSRC (BB/N017773/2), SNF (CRSK-3_190550; IZSEZ0_204655), Vontobel (41309), and the UZH-URPP (Translational Cancer Research). KY is supported by a BioMedTech Entrepreneur Fellowship.

## AUTHOR CONTRIBUTIONS

OK, KY and RC conceived the idea and coordinated the project. RC secured funding and guided the work. OK performed experiments and analyses. AA performed nFCM vs TEM sizing comparison. BP provided support on nFCM analysis and markers panel design. OK and KY performed statistical data analyses. OK and KY wrote the manuscript draft, which was edited by RC.

## SUPPLEMENTAL MATERIAL

All supplementary material are included in a single combined file in PDF format.

