## Supplementary data for "Assessing extracellular vesicles in human biofluids using flow-based analyzers"

### Figure S1

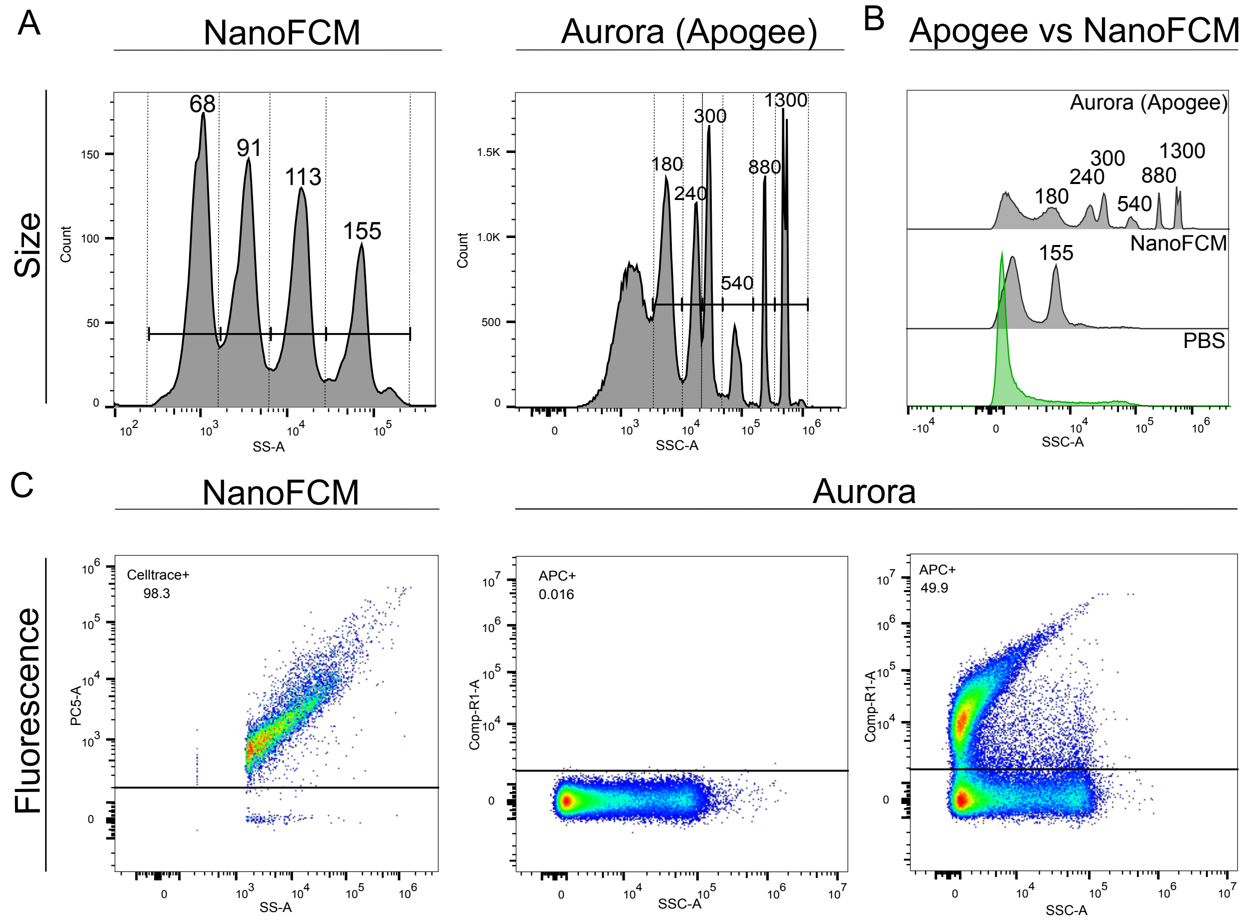


**Figure S1.** Representative scatter histogram showing size binning strategy (top) and fluorescence gating strategy (bottom).

### Figure S2

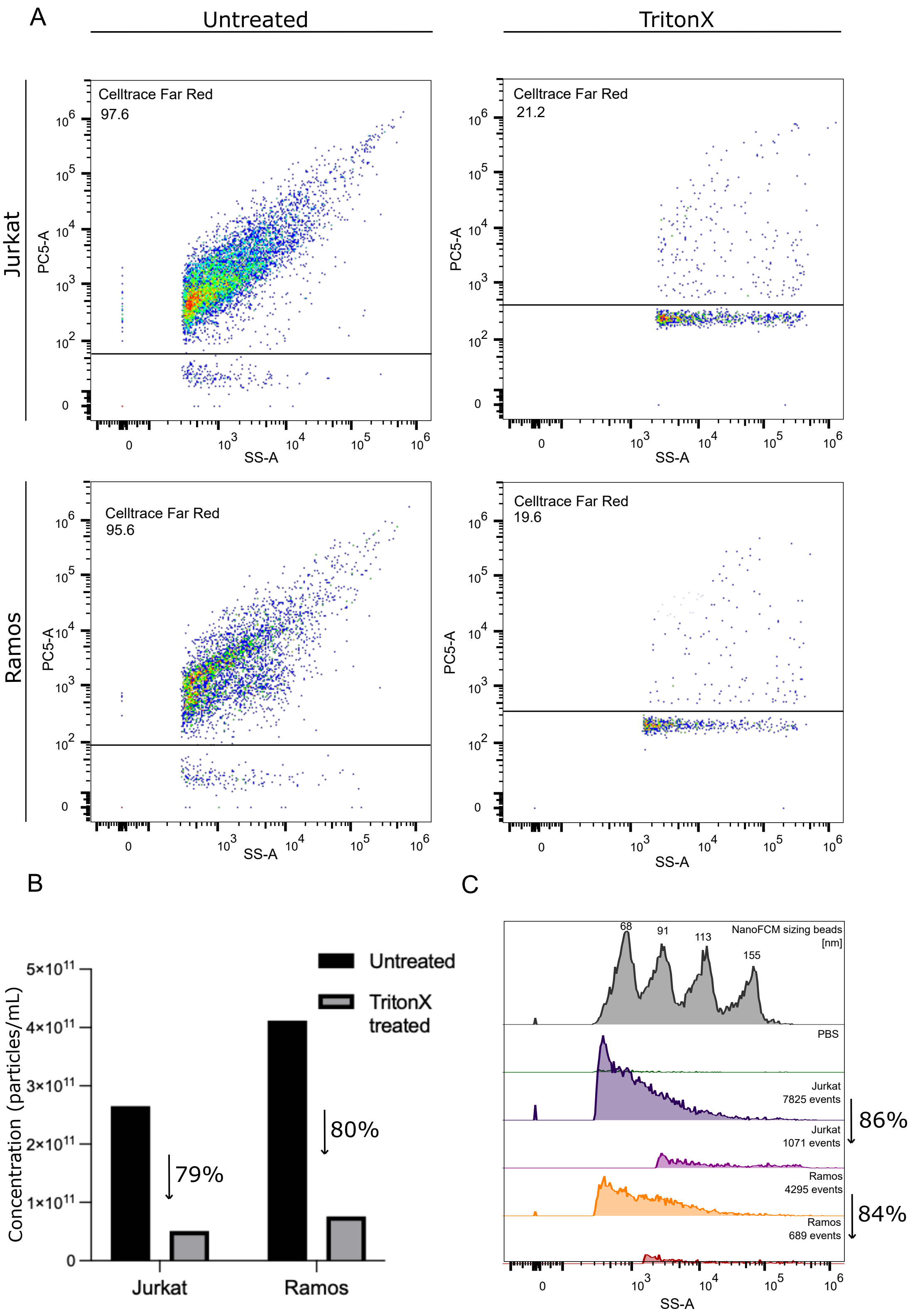


**Figure S2.** Purity assessment of Jurkat and Ramos cell-line derived sEVs (n = 3). (A) Representative figures showing reduction of detectable celltrace fluorescence before and after Cell-line EV treatment with 1% TritonX detergent. (B) Bar graph showing the reduction in mean particle concentration in EV samples treated with the detergent. Percentages indicate the reduction in concentration after treatment (C) Representative side scatter histogram of nfcm sizing beads, PBS as well as Jurkat and Ramos EVs before and after treatment with detergent. Percentages indicate the reduction in concentration after treatment.

### Figure S3

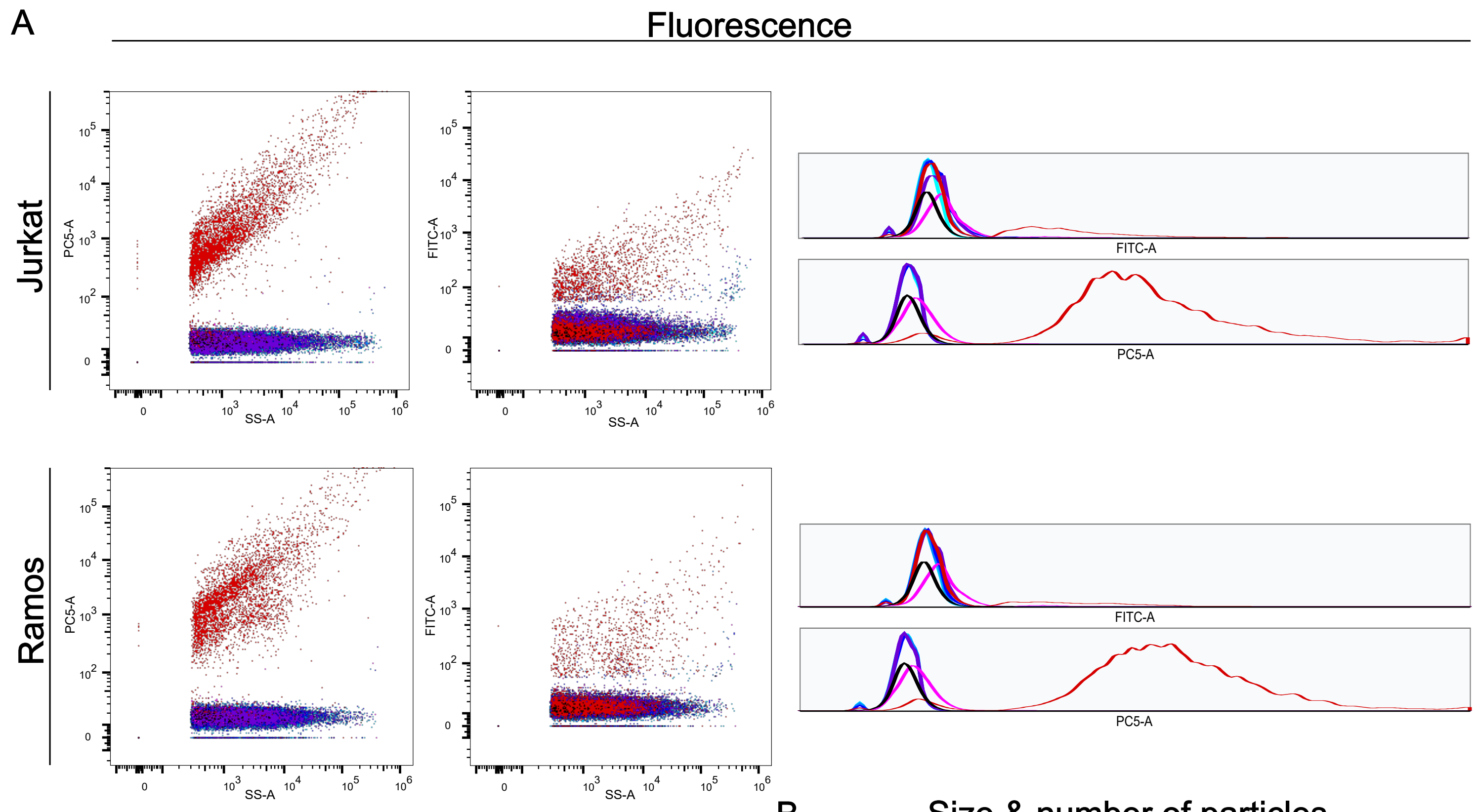


| | Sample |
|---|---|
| ■ (black) | PBS |
| ■ (red) | EV, Celltrace - CD63(FITC) |
| ■ (magenta) | Antibody only, CD63 (APC) - CD45 (FITC) |
| ■ (purple) | EV, IgG2a (rat) (FITC) |
| ■ (blue) | EV, IgG2a (mouse) (FITC) |
| ■ (light blue) | EV, IgG1(mouse) (FITC) |
| ■ (cyan) | EV (unstained) |

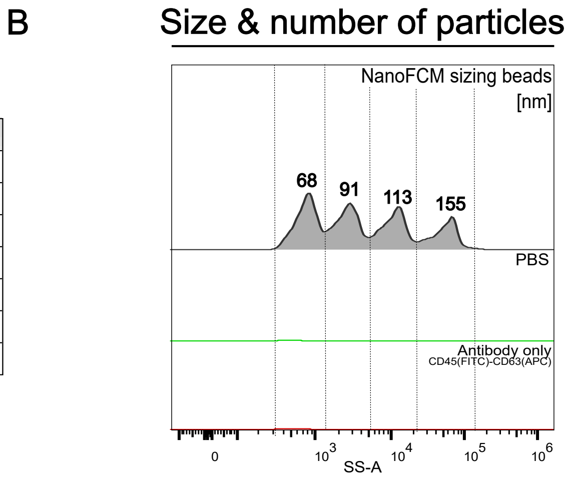


**Figure S3**. Negative controls used in cell-line EVs experiments. (A) Representative scatter plots and histograms showing fluorescence detection of samples including filtered PBS, cell-line EVs stained with CellTrace far red and CD63 (FITC), antibody only (no EVs), EVs stained with isotype controls (IgG2a (mouse and rat) and IgG1 (mouse)) as well as unstained EVs. (B) Size detection of PBS and antibody only samples.

**Figure S4**

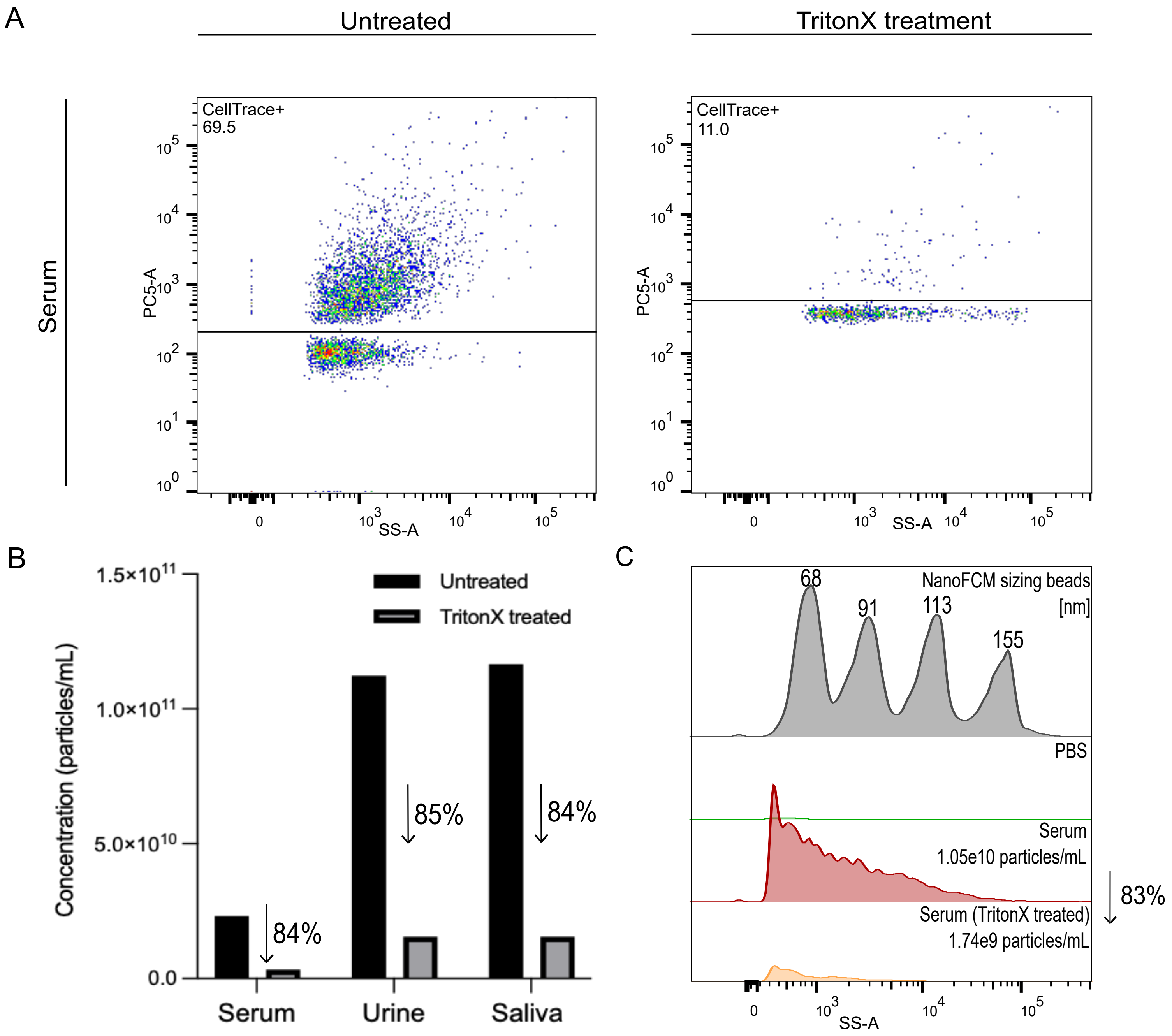


**Figure S4.** Purity assessment of purified sample serum sEVs. (A) Representative figures showing reduction of detectable celltrace fluorescence before and after serum EV treatment with 1% TritonX detergent. (B) Bar graph showing the reduction in mean particle concentration in EV samples treated with the detergent. Percentage indicates the reduction in concentration after treatment. (C) Representative side scatter histogram of nfcm sizing beads, PBS as well as serum EVs before and after treatment with detergent. Percentage indicates the reduction in concentration after treatment.

**Figure S5**

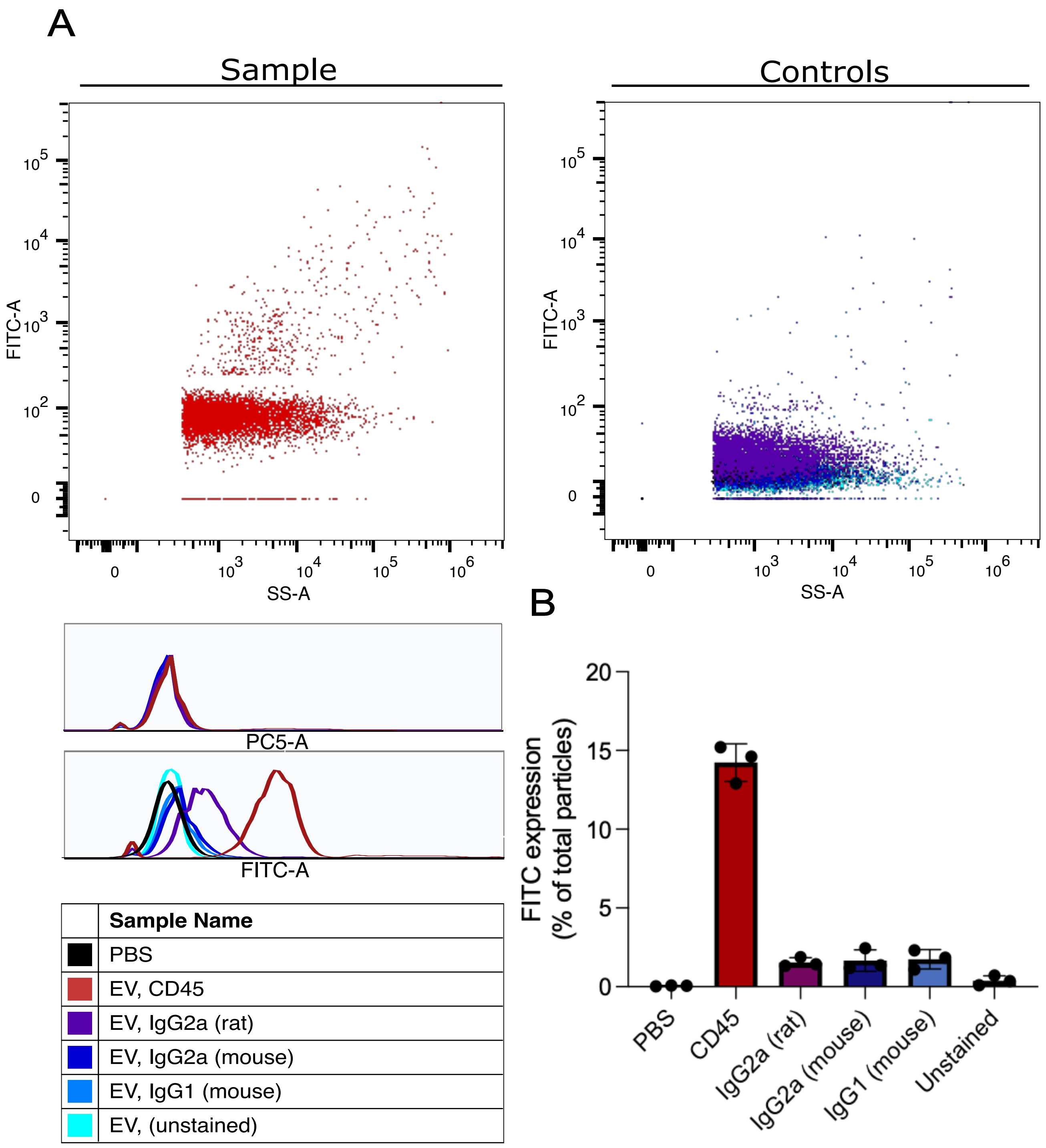


**Figure S5**. Negative controls used in serum EVs experiments. (A) Representative scatter plots and histograms showing fluorescence detection of samples including filtered PBS, serum EVs stained with CD45 (FITC), antibody only (no EVs), EVs stained with isotype controls (IgG2a (mouse and rat) and IgG1 (mouse)) as well as unstained EVs. (B) Size detection of PBS and antibody only samples.

### Figure S6

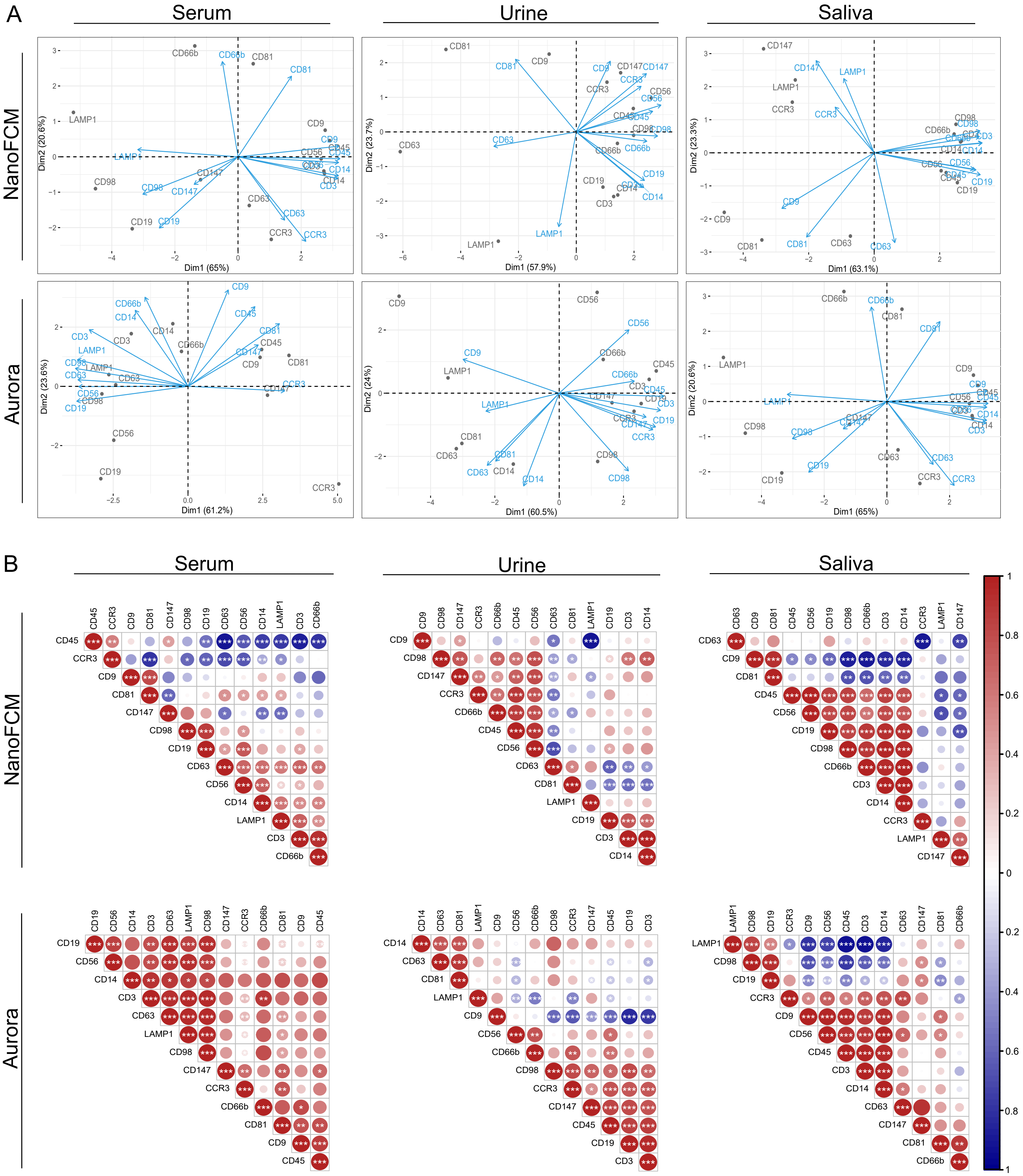


**Figure S6**. PCA and correlation of biofluids derived sEVs markers measured in NanoFCM® nanoflow analyzer and Cytek® Aurora. (A) PCA bi-plot analysis of EVs markers measured by NanoFCM and Aurora in different sources of biofluids EVs. (B) Spearman's rank correlation matrix of EVs markers measured in NanoFCM and Aurora from serum, urine and saliva EVs. N=4. One-way ANOVA, $p < 0.05$ *, $p < 0.01$ **, $p < 0.005$ ***.

**Table S1.** Antibodies and reagents list.

| Marker | Fluorophore | Reactivity | Clone | Company | Cat. No. | Isotype |
|---|---|---|---|---|---|---|
| CCR3 | APC | Human | 5E8 | Biolegend | 310707 | Mouse IgG2b, κ |
| CD3 | FITC | Human | HIT3a | Biolegend | 300305 | Mouse IgG2a, κ |
| CD9 | APC | Human | HI9a | Biolegend | 312107 | Mouse IgG1, κ |
| CD9 | FITC | Human | HI9a | Biolegend | 312103 | Mouse IgG1, κ |
| CD14 | FITC | Human | M5E2 | Biolegend | 301803 | Mouse IgG2a, κ |
| CD19 | APC | Human | HIB19 | Biolegend | 302211 | Mouse IgG1, κ |
| CD19 | FITC | Human | HIB19 | Biolegend | 302205 | Mouse IgG1, κ |
| CD45 | APC | Human | HI30 | Biolegend | 304011 | Mouse IgG1, κ |
| CD45 | FITC | Human | HI30 | Biolegend | 304006 | Mouse IgG1, κ |
| CD56 | FITC | Human | HCD56 | Biolegend | 318303 | Mouse IgG1, κ |
| CD63 | APC | Human | H5C6 | Biolegend | 353007 | Mouse IgG1, κ |
| CD63 | FITC | Human | H5C6 | Biolegend | 353005 | Mouse IgG1, κ |
| CD66b | FITC | Human | G10F5 | Biolegend | 305103 | Mouse IgM, κ |
| CD81 | APC | Human | 5A6 | Biolegend | 349509 | Mouse IgG1, κ |
| CD81 | FITC | Human | 5A6 | Biolegend | 349503 | Mouse IgG1, κ |
| CD98 | FITC | Human | MEM-108 | Biolegend | 315603 | Mouse IgG1, κ |
| CD147 | APC | Human | HIM6 | Biolegend | 306213 | Mouse IgG1, κ |
| CellTrace Far Red | APC | | | Thermo | C34572 | n/a |
| LAMP1 | FITC | Human | H4A3 | Biolegend | 328605 | Mouse IgG1, κ |
